# A Nonsteroidal Reversal Agent Inhibits Allopregnanolone Modulation of α1β3δ GABA_A_ Receptors

**DOI:** 10.64898/2026.04.14.718525

**Authors:** Xiaojuan Zhou, Youssef Jounaidi, Keith W. Miller

**Affiliations:** Department of Anesthesia, Critical Care and Pain Medicine, Harvard Medical School, Massachusetts General Hospital, 32 Fruit Street, Boston, Massachusetts 02114, United States; Department of Anesthesia, Critical Care and Pain Medicine, Harvard Medical School, Massachusetts General Hospital; Department of Anesthesia, Critical Care and Pain Medicine, Massachusetts General Hospital

**Keywords:** GABA_A_R, Allopregnanolone, Reversal Agent, Spiro-hydantoin, Allosterism

## Abstract

The neurosteroid allopregnanolone is a positive allosteric modulator of GABA(A) receptors, which has proved beneficial in the treatment of major depressive disorder and epilepsies. It also has a role in treating the mood swings that are associated with fluctuations in its level during the menstrual cycle. Nonetheless, a subset of women do not tolerate high levels of allopregnanolone. Iso-allopregnanolone, a negative allosteric modulator, as well as synthetic steroid antagonists are used to treat such conditions. However, steroid-based medications are difficult to deliver and their specificity of action can be unclear. Recently introduced novel nonsteroidal agents that, like iso-allopregnanolone, can reverse the action of positive allosteric modulators without changing the positive action of GABA, might provide an alternative. We surveyed a number of them on human α1β3δ GABA_A_Rs using a [^3^H]muscimol binding assay. A 6-membered ring spiro-hydantoin, DKD99, allosterically reversed the positive allosteric action of allopregnanolone over a wide concentration range (6 to 1,000 nM). DKD99 shifted allopregnanolone’s modulation curve 10-fold to the right. Furthermore, it has a much lower affinity when exerting similar actions on α1β3γ2 receptors. Agents such as this have utility for elucidating underlying mechanisms and may offer an alternative pathway for the development of nonsteroidal therapies against the positive allosteric modulatory actions of neurosteroids.

## INTRODUCTION

Allopregnanolone (5α-Pregnan-3α-ol-20-one) is an endogenous neurosteroid that regulates affective disorders via its positive allosteric modulator actions on GABA_A_Rs.^1^ It has attracted the attention of pharmacologists because of its therapeutic potential in treating depression and epilepsy.^2-5^ This has further lead to the development of Brexanalone, a formulation of allopregnanolone for intravenous delivery and Zuranalone, a steroid that can be taken orally.^6, 7^

However, the physiological concentration of allopregnanolone fluctuates in response to various inputs and this can affect mood. Nowhere is this more apparent than in the menstrual cycle. In post-menstrual dysphoric disorder (PMDD) the level of allopregnanolone fluctuates throughout the cycle and is associated with mood swings, irritability, anxiety and depression. ^2, 8-10^ In PMDD the role of fluctuating allopregnanolone levels and their link to behavior is complex, with levels rising by an order of magnitude between the follicular and luteal phases. It is then paradoxical that in a subset of women PMDD is experienced during this phase. However, the neurosteroid iso-allopregnanolone (5β-Pregnan-3α-ol-20-one), an endogenous negative allosteric modulator, also fluctuates during the cycle.^11, 12^ Although the mechanism remains unclear and may involve fluctuations in the isoforms of the GABA_A_R involving the δ-subunit, this has led to the introduction of negative allosteric modulator therapeutics such as Sepranolone and Golexanalone (GR-3027).^1, 5, 7, 12-14^ Research with such GABA_A_ receptor modulating steroid antagonists (GAMSAs) has revealed that they may also have a therapeutic role to play in diverse etiologies, often related to neuro-inflammation, such as cognitive function, hepatic encephalopathy and motor incoordination.^15-17^

Neurosteroids act on most isoforms of the GABA_A_R but are particularly effective on δ-subunit containing receptors partly because GABA is a partial agonist and the potential for enhancing action is corresponding high.^18,19^ Studies of δ-subunit containing extrasynaptic receptors in heterologous systems have been bedeviled by promiscuous assembly in α4/6βxδ receptors. ^20-23^ However, the α1βxδ subunits are thought to assemble homogeneously as in α1βγ2 receptors (β–α1–δ–β–α1) and were therefore chosen for this study. ^24-26^

Recently, a series of spiro-barbiturates and spiro-hydantoins have been assayed for their ability to reverse the action of positive allosteric modulators on synaptic (α1β3γ2) and extra-synaptic (α1β3) GABA_A_Rs. Some of these agents acted negative allosteric modulators of positive allosteric modulators including the steroid general anesthetic alphaxalone but. as null allosteric ligands of orthosteric agonist binding. They have been termed reversal agents and some progress has been made towards discovering their mechanism of action.^27-30^ Their actions resemble those reported for iso-allopregnanolone ^12, 31^. In this report, we have evaluated a subset of these compounds on heterologously expressed human full length α1β3δ GABA_A_Rs (**Fig. 1**). Although, the structure activity relationships differed from their action on α1β3γ2 receptors, two of them selectively reversed allopregnanolone’s positive allosteric modulation at low micromolar concentrations that had no action on α1β3γ2 receptors.

**Figure 1.**
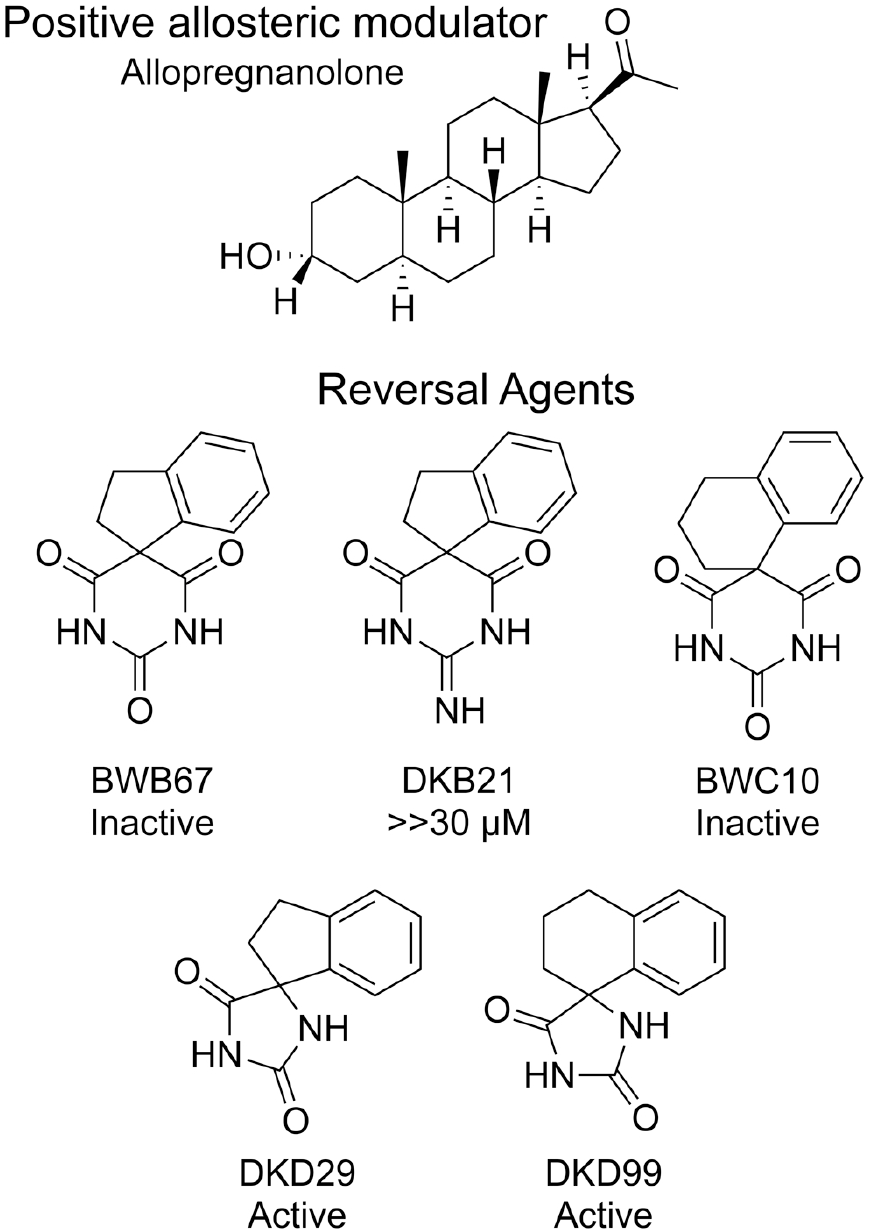
The chemical structures of the ligands studied. Top row: Allopregnanolone or 3α-Hydroxy-5α-pregnan-20-one. Middle row: Spiro-barbiturate reversal agents. Bottom row: Spiro-hydantoin reversal agents.

## RESULTS

### Principle of the assay

We assayed for reversal activity at equilibrium using an [^3^H]muscimol binding assay based on the observation that the GABA_A_R exists in a dynamic equilibrium between a low affinity resting state and a smaller fraction of receptors in a high affinity desensitized state.^32^ By using a low concentration of [^3^H]muscimol that mainly binds to the small fraction of high affinity receptors, the action of positive allosteric modulators (PAMs), such as neurosteroids and general anesthetics, to stabilize the desensitized state can be observed. The assay is a convenient way to test for agents that reverse positive allosteric modulator action. It has been employed in αβγ and αβ receptors with success, but this is the first time it has been employed in α1β3δ receptors.

**Table 1.** Reversal agents have little action on [^3^H]muscimol binding.

| Agent | Average ± SD | N |
| --- | --- | --- |
| DKD21 | 100 ± 0.02 | 15 |
| BWC10 | 118 ± 12.7 | 17 |
| DKD29 | 99 ± 5.1 | 23 |
| DKD99 | 103 ± 4.9 | 27 |
| Control | = 100 |  |

### Do reversal agents interact with the orthosteric agonist site

It is important to ensure that reversal agents do not displace [^3^H]muscimol from its binding site. Each reversal agent in **Fig. 1** was titrated between 0.1 and 100 μM against 3 nM [^3^H]muscimol binding in α1β3δ GABA_A_Rs. None decreased [^3^H]muscimol binding and one, BWC10, was a weak PAM, enhancing [^3^H]muscimol from 100 to 125 ± 4.5%. This is of little functional significance compared to the maximum enhancement caused by 100 μM etomidate of 441 ± 19% (n = 22).

### The structural dependence of reversal action

We tested the ability of the reversal agents in **Fig. 1**, to reverse the positive allosteric action of 100 nM allopregnanolone on α1β3δ GABA_A_Rs with the specific goal of discovering a reversal agent with good efficacy, and an IC50 in the low micromolar range. We chose 100 nM allopregnanolone for this survey because it enhanced [^3^H]binding by 288 ± 13% (n = 6) making it easier to detect reversal activity.

None of the three spiro barbiturates reversed allopregnanolone’s enhancing action with the exception of DKB21, which was inactive at 30 μM but at 100 μM modestly reduced allopregnanolone’s enhancing action by 10 % (n = 5, p= 0.02).

In contrast, both of the spiro-hydantoins tested reversed allopregnanolone’s action. The 5-membered ring hydantoin, DKD29, modestly reversed allopregnanolone-enhanced [^3^H]muscimol binding from 325 to 253% without displacing [^3^H]muscimol binding itself (**Fig. 2**), whereas the 6-membered ring hydantoin, DKD99, was twice as efficacious. Although efficacy was dependent on ring size, potency was not, and both agents had similar IC50s of 5 μM (**Table 2**). Encouragingly, this compares to IC50s of 40 μM in synaptic α1β3γ2 GABA_A_Rs indicating that this reversal action is quite selective for extrasynaptic α1β3δ receptors.

**Table 2.** Reversal curves for allopregnanolone’s enhancement of [^3^H]muscimol binding.

| Reversal agent | 3α5αP nM | 1C50 μM | ± | SD | Max | ± | SD | Min | ± | SD | Normalized Efficacy | ± | SD |
| --- | --- | --- | --- | --- | --- | --- | --- | --- | --- | --- | --- | --- | --- |
| DKD99 | 6 | 2.2 | ± | 0.3 | 219 | ± | 2.7 | 92 | ± | 3.1 | 0.58 | ± | 0.01 |
| DKD99 | 100 | 5.0 | ± | 0.6 | 264 | ± | 2.3 | 115 | ± | 3.6 | 0.56 | ± | 0.01 |
| DKD99 | 1,000 | 5.2 | ± | 1.7 | 331 | ± | 5.3 | 214 | ± | 8.6 | 0.35 | ± | 0.01 |
| DKD29 | 100 | 4.5 | ± | 1.1 | 327 | ± | 2.6 | 253 | ± | 4.0 | 0.23 | ± | 0.01 |

Normalized efficacy is the difference between the maximum and minimum enhancement normalized to the maximum enhancement.

**Figure 2.**
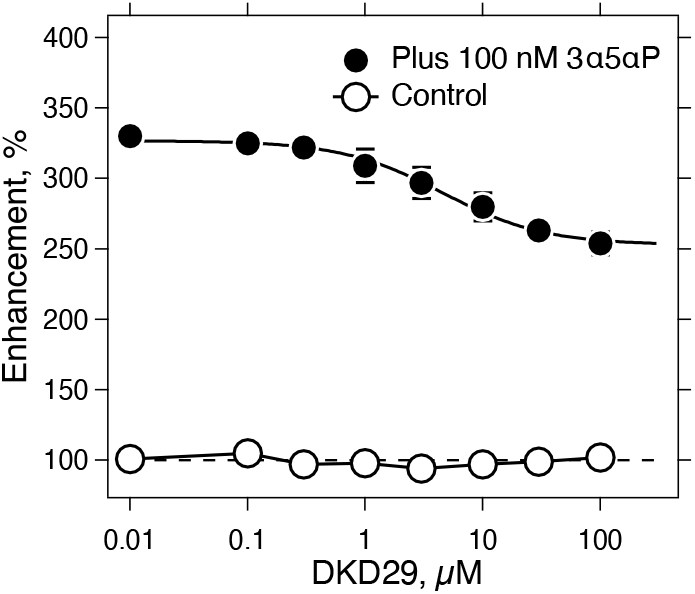
DKD29 partially reverses allopregnanolone’s PAM action on α1β3δ GABA_A_Rs without displacing [^3^H]muscimol binding. N =3 at each concentration. Standard deviations are shown when larger than symbols.

Because of its higher efficacy we studied DKD99 in more detail. Allopregnanolone’s enhancement of [^3^H]muscimol binding was reversed by DKD99 at 6, 100 and 1,000 nM. At both 6 and 100 nM allopregnanolone reversal was close to complete, but at 1 μM allopregnanolone reversal was far from complete perhaps indicating a ceiling effect in the allosteric interaction between the allopregnanolone sites and the reversal site(s) (**Table 2, Fig. 3**).

**Figure 3.**
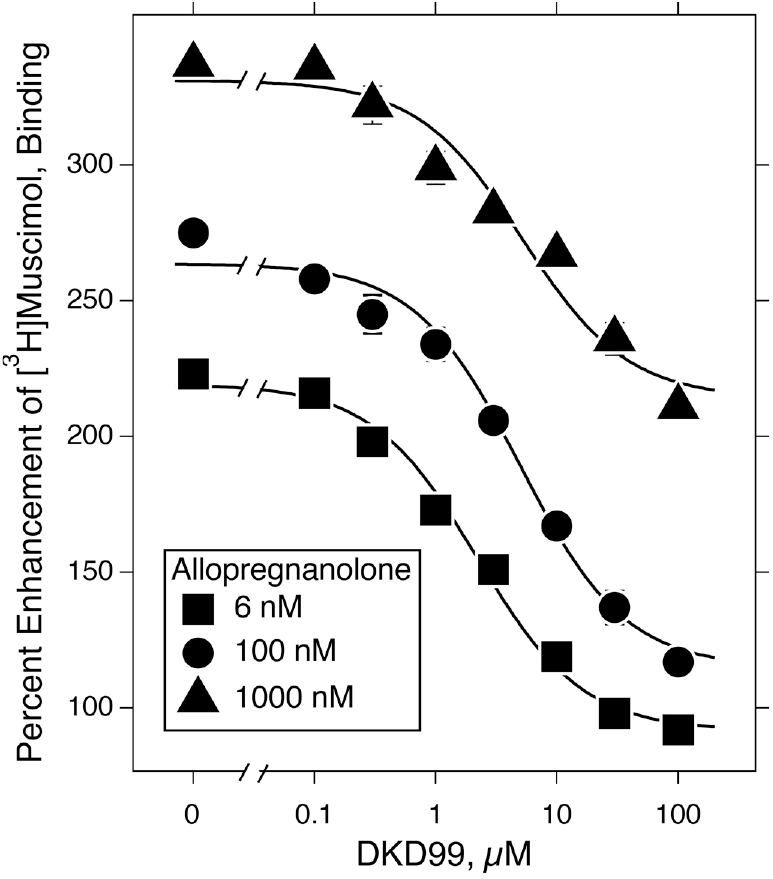
DKD99 reverses allopregnanolone’s enhancement of [^3^H]muscimol binding over a wide concentration range. Each point determined in triplicate.

### Allopregnanolone enhances [^3^H]muscimol binding over a wide range of concentrations

Allopregnanolone enhanced [^3^H]muscimol binding in a concentration-dependent manner with enhancement reaching 20% between 0.1 and 0.3 nM and plateauing at 370% at ≥10 μM, which is 86% of that for etomidate (**Fig. 4**). Etomidate and steroids both bind in the same β^+^/α– interfaces in the transmembrane domain but etomidate is situated closer to the orthosteric agonist site than steroids. This may offer an explanation for the difference in enhancing efficacy. The enhancement curve had a midpoint of 14 nM and a Hill coefficient of 0.5 (**Table 3**).

**Table 3.** The concentration-dependence of allopregnanolone’s enhancement of [^3^H]muscimol binding in α1β3δ GABA_A_Rs in the absence and presence of 50 μM of the reversal agent DKD99.

| Equation | Parameter | $\alpha 1\beta 3\delta$ | ± | SD | | N | $\alpha 1\beta 3\delta$<br>+<br>50 μM<br>DKB99 | ± | SD | | N |
| --- | --- | --- | --- | --- | --- | --- | --- | --- | --- | --- | --- |
| Hill | EC50 | 14 | ± | 1.9 | nM | 78 | 141 | ± | 51 | nM | 52 |
| | nH | 0.51 | ± | 0.03 | | | 0.46 | ± | 0.05 | | |
| | Max | 378 | ± | 5.1 | | | 287 | ± | 10 | % | |
| Adair | K1 | 1.3 | ± | 0.23 | nM | 78 | 0.85 | ± | 0.29 | nM | 52 |
| | K2 | 148 | ± | 32 | nM | | 629 | ± | 106 | nM | |
| | Fraction Site 1 | 0.52 | ± | 0.03 | | | 0.28 | ± | 0.02 | | |
| | Max | 371 | ± | 3.3 | % | | 285 | ± | 3.8 | % | |

The concentration of [<sup>3</sup>H]muscimol was 3 nM. Each curve is the combined data from four experiments.

**Figure 4.**
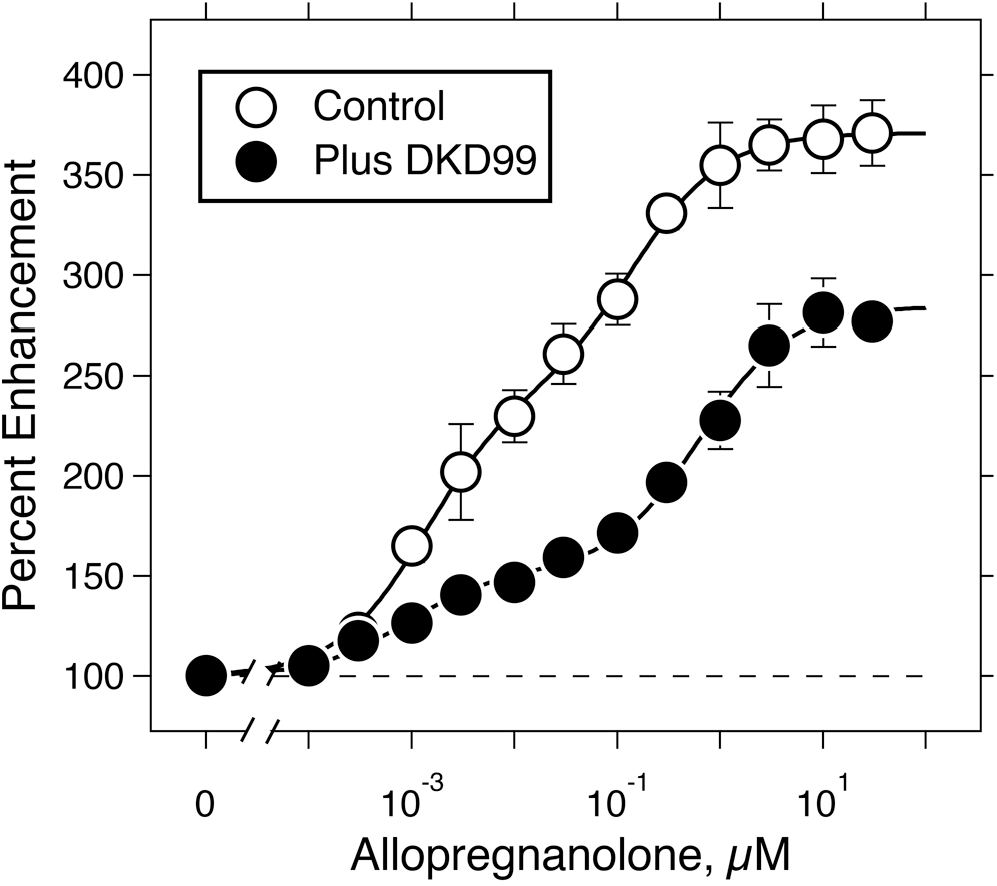
In α1β3δ GABA_A_Rs, the concentration-dependence allopregnanolone’s enhancement of [^3^H]muscimol binding shifted to the right and reduced in amplitude in the presence of the reversal agent DKD99 (50 μM). The curves are fits to a two site Adair equation. Number of data points: Control, 89; +DKD99, 53.

### DKD99 shifts allopregnanolone’s enhancement curve to the right

Reversal agents that act allosterically are expected to shift this curve to the right. ^27^ To test this we repeated the above titration in the presence of a fixed concentration (50 μM) of the spiro-hydantoin DKD99. DKD99 reversed allopregnanolone’s action over the whole concentration range (6 to 1,000 nM). Compared to the control curve, DKD99 shifted allopregnanolone’s enhancement curve 10-fold to the right and lowered its maximum some 30% without changing its Hill coefficient. The low Hill coefficient suggests that there is more than one site or process underlying the enhancement.

### Interpretation of the low Hill coefficient

The goal of this study was to test whether reversal agents can reverse the action of allopregnanolone on α1β3δ receptors so we did not seek the cause of the low Hill coefficient. It has been claimed that in α1β2γ2 receptors there are three noninteracting steroid sites that act independently.^33^However, a single particle Cryo–EM structure of α1β2γ2 receptors in the presence of GABA and allopregnanolone shows it bound to the two classic sites in the β^+^/α^−^ interface in the transmembrane domain. ^34^ In the absence of a structure for the α1β3δ receptor, the low Hill coefficient could be interpreted in several ways. If there are two or more different allopregnanolone sites, they could either act independently but have different affinities or they could have similar affinities but interact with negative allosterism. Alternatively, there could be two different states or conformations that have high affinity for [^3^H]muscimol but different affinities for allopregnanolone. We will call the two actions “components” to avoid implying a mechanism.

We chose simply to fit the data to a two independent binding site model to provide a robust description that aims to deconvolute the two phases of action. This model is sometimes referred to as a two site Adair equation.

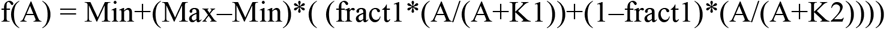

where Max and Min are the respective amplitudes at zero and the plateau allopregnanolone concentration, and A is the concentration of allopregnanolone. K1 and K2 are the dissociation constants of the two sites or conformations and fract1 is the fractional population of site or conformation 1.

This analysis deconvoluted allopregnanolone’s concentration-response curve into a high affinity component with a dissociation constant of 1 nM and a low affinity component whose dissociation constant was some 100-fold higher. The population of these two components was distributed equally (**Table 2**).

Fitting the data to a two site Adair equation revealed that the high and low affinity sites reacted to DKD99 differently, which supports the idea that they represent different states. DKD99 acted on allopregnanolone’s high affinity site to decrease its fractional contribution without changing its EC50. In contrast, DKD99 shifted allopregnanolone’s low affinity site’s EC50 4-fold to higher concentrations without changing its overall contribution to enhancement. That is, the decrease in overall enhancement originates entirely from DKD99’s action on allopregnanolone’s high affinity site, and the right shift from its low affinity site. This conclusion, unlike the Hill equation’s description, is model-dependent and determining the mechanism of these unexpected functional actions of DKD99 will require more detailed work including structural studies.

## CONCLUSION

The spiro-hydantoin reversal agent, DKD99, reverses with micromolar potency and good efficacy allopregnanolone’s positive allosteric action on extrasynaptic α1β3δ GABA_A_Rs over a wide concentration range. Previously, reversal of allopregnanolone’s positive allosteric actions has only been accomplished with steroid antagonists that can be challenging to formulate for clinical use and whose metabolism may have downstream effects. Nonsteroidal agents such as DKD99 may both prove useful in dissecting mechanisms in more complex systems and provide a novel starting point for therapeutic development.

## EXPERIMENTAL METHODS

### Cell line creation

A new α1β3δ inducible HEK293 cell line was created using constructs for human full length β3- and δ-subunits that have been previously described ^35-37^. A new full length human α1-subunit (Uniprot P14867) construct was synthesized (Synbio technologies, Inc) bearing an N-terminal twin strep tag (YPYDVPDYAGGSWSHPQFEKGGGSGGGSGGSAWSHPQFEK) inserted five amino acid after the α1-subunit signal peptide cleavage site and followed by a flexible linker GGS. The α1-twin strep tagged subunit was cloned into Doxycycline inducible expression plasmid pCDNA4TO (EcoRI-XhoI), with selection marker Zeocin (See Supplemental Materials).

The α1-, β3- and δ-constructs were stably transfected into a HEK293T TetR cell line with Lipofectamine 2000 reagent as directed by the manufacturer’s protocol. The transfection ratio of α1- and δ-subunits being 2 : 0.25 and using ratios for the β3-subunit of either 0.3 or 1. At 48 hours post-transfection cells were trypsinized and plated in the presence of Zeocin 250 μg/ml, Hygromycin 50 μg/ml and G418 100u μg/ml with 5 μg/ml Blasticidin for one week. The selected pools were then sorted twice by flow cytometry for GABA receptor expression using APC conjugated anti-1D4 antibodies against the 1D4-tagged delta subunit as a surrogate for a full pentameric expression at the cell surface. The twice enriched pool was then used to isolate clones by single cell plating in a 96 well plate and antibiotic selection for another week. The 18 best growing clones were then analyzed by flow cytometry using the same methodology and the best clones were further characterized by [^3^H]muscimol binding assay.

### Selection and pharmacological characterization of clones

Judged by FACS the lower ratio of β3-subunit gave better yields overall, but individual clones from both transfection ratios gave good specific activities. The clone selected was from the 2:0.3:0.25 (α1:β3:δ) group. It had a good growth rate and a specific activity of 17 ± 2 pmol of muscimol binding sites/mg of membrane protein, comparable to a similar cell line expressing α1β3γ2 receptors.^36^ DS2 is a δ-subunit selective ligand.^38^ It strongly enhanced [^3^H]muscimol binding by 315 ± 25% in α1β3δ receptors compared to 8 ± 3% in α1β3γ2 receptors. We estimated the dissociation constant of GABA by titrating it against 3 nM [^3^H]muscimol, a concentration chosen as a compromise between low occupancy and having sufficient cpm. The titration from 1 nM to 1 mM yielded an IC50 of 150 ± 10 nM (n = 32; 3 separate experiments). A similar experiment with α1β3γ2 receptors yielded an IC50 of 75 ± 9 μM (n = 34; 2 separate experiments). The difference in IC50s is consistent with the higher agonist affinity of α1β3δ over α1β3γ2receptors.

### The [^3^H]muscimol binding assay

The methods have been recently described.^27^ Briefly, binding assays were performed on cell membranes harvested from HEK cells. [^3^H]muscimol (PerkinElmer) was used at 3 nM final concentration and nonspecific binding was corrected for by displacement of specific binding by 1 mM GABA. Samples were made in 7 mL glass sample vials. Allopregnanolone was added to glass sample vials after serial dilution from a 20 mM stock in DMSO. Samples were equilibrated for 30 min and filtered on GFB filters, which were dried under a lamp before being added to 5 mL of scintillation cocktail and counted. Analysis was carried out using Wavemetrics’s Igor software.

## Acknowledgements

This research was supported by a grant from the National Institutes of Health, USA, R01 GM135550, and with support from the Department of Anesthesia, Critical Care and Pain Medicine, Massachusetts General Hospital, Boston.

## Supplemental Materials

### Human α1 N-Terminal Twin-strep based on Uniprot # P14867

Red is the signal peptide.

Green is the HA tag 9-amino acid peptide epitope (YPYDVPDYA) derived from the human influenza hemagglutinin (HA) molecule.

Blue is the two Twin Strep tags (WSHPQFEK) connected by a flexible linker.

Yellow are flexible linkers.

### Protein sequence

MRKSPGLSDCLWAWILLLSTLTGRSYGQPSLQYPYDVPDYAGGSWSHPQFEKGGGSGGGSGGSAWSHPQFEKGGSDELKDNTTVFTRILDRLLDGYDNRLRPGLGERVTEVKTDIFVTSFGPVSDHDMEYTIDVFFRQSWKDERLKFKGPMTVLRLNNLMASKIWTPDTFFHNGKKSVAHNMTMPNKLLRITEDGTLLYTMRLTVRAECPMHLEDFPMDAHACPLKFGSYAYTRAEVVYEWTREPARSVVVAEDGSRLNQYDLLGQTVDSGIVQSSTGEYVVMTTHFHLKRKIGYFVIQTYLPCIMTVILSQVSFWLNRESVPARTVFGVTTVLTMTTLSISARNSLPKVAYATAMDWFIAVCYAFVFSALIEFATVNYFTKRGYAWDGKSVVPEKPKKVKDPLIKKNNTYAPTATSYTPNLARGDPGLATIAKSATIEPKEVKPETKPPEPKKTFNSVSKIDRLSRIAFPLLFGIFNLVYWATYLNREPQLKAPTPHQ*

### Nucleotide sequence

ATGAGAAAGAGCCCTGGCCTGAGCGATTGTCTGTGGGCCTGGATTCTGCTGCTGAGCACCCTGACAGGCAGAAGCTATGGCCAGCCTAGCCTGCAGTACCCCTACGACGTGCCAGATTATGCCGGCGGATCTTGGAGCCATCCTCAGTTCGAAAAAGGCGGCGGTTCTGGCGGTGGATCTGGCGGATCTGCTTGGTCACACCCACAGTTTGAGAAAGGCGGAAGCGACGAGCTGAAGGACAACACCACCGTGTTCACCAGAATCCTGGACAGACTGCTGGACGGCTACGACAACAGACTGAGGCCTGGCCTCGGCGAGAGAGTGACCGAAGTCAAGACCGACATCTTCGTGACCAGCTTCGGCCCCGTGTCCGACCACGATATGGAGTACACCATCGACGTGTTCTTCCGGCAGAGCTGGAAGGACGAGCGGCTGAAGTTTAAGGGCCCCATGACCGTGCTGCGGCTGAACAATCTGATGGCCAGCAAGATCTGGACCCCTGACACATTCTTCCACAACGGCAAGAAAAGCGTGGCCCACAACATGACCATGCCTAACAAGCTGCTGCGGATCACCGAGGATGGCACCCTGCTGTACACCATGAGGCTGACAGTCAGAGCCGAGTGTCCCATGCACCTGGAAGATTTCCCTATGGACGCCCACGCCTGTCCTCTGAAGTTTGGCAGCTACGCCTACACAAGAGCCGAGGTGGTGTACGAGTGGACCAGAGAACCTGCCAGATCTGTGGTGGTGGCCGAGGACGGAAGCAGACTGAACCAGTATGATCTGCTGGGCCAGACCGTGGACTCTGGCATTGTGCAAAGCAGCACCGGCGAGTACGTGGTCATGACAACCCACTTCCACCTGAAGCGGAAGATCGGCTACTTCGTGATCCAGACCTACCTGCCTTGCATCATGACAGTGATCCTGAGCCAGGTGTCCTTCTGGCTGAACCGGGAATCTGTGCCTGCCAGAACAGTGTTCGGCGTGACCACCGTGCTGACCATGACCACACTGAGCATCAGCGCCAGAAACAGCCTGCCTAAGGTGGCCTACGCCACCGCTATGGACTGGTTTATC

GCCGTGTGCTACGCCTTCGTGTTCAGCGCCCTGATCGAGTTCGCCACCGTGAACTACTTCACCAAGAGAGGCTACGCCTGGGACGGCAAGTCTGTGGTGCCAGAGAAGCCCAAGAAAGTGAAGGACCCTCTGATCAAGAAGAACAACACATACGCCCCTACCGCCACCAGCTACACCCCTAATCTTGCCAGAGGCGATCCTGGCCTGGCCACAATTGCCAAGTCTGCCACCATCGAGCCTAAAGAAGTGAAGCCCGAGACAAAGCCTCCTGAGCCTAAGAAAACCTTCAACAGCGTGTCCAAGATCGACCGGCTGAGCCGGATTGCCTTTCCTCTGCTGTTCGGCATCTTCAACCTGGTGTACTGGGCCACCTACCTGAACAGAGAGCCCCAGCTGAAAGCCCCTACACCTCACCAGTGA

### Clone selection

**Figure S1.**
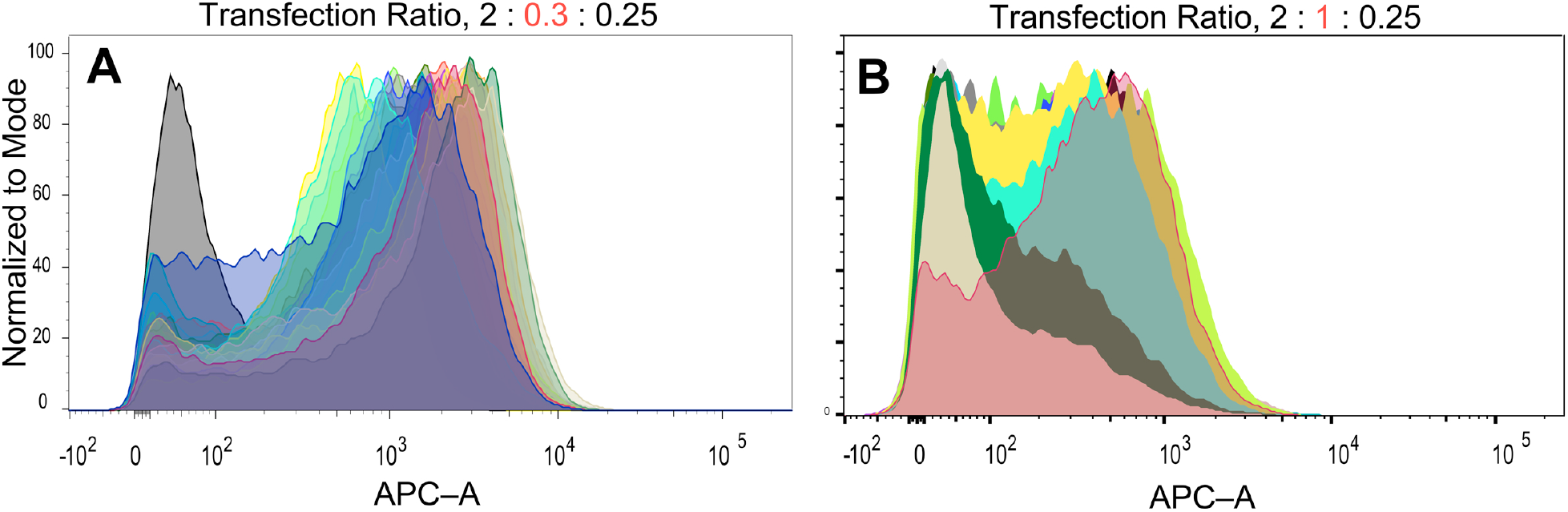
Flow cytometry selection analysis of α1-, β3- and δ-constructs stably transfected into a HEK293T TetR clonal cell lines using two transfection ratios α1:β3:δ: **A**, 2:0.3:0.26 or **B**, 2:1:0.25. The isolated clones were sorted for GABA_A_R receptor expression using APC conjugated anti-1D4 antibodies against the 1D4-tagged delta subunit. Cells were analyzed at the MGH Flow Cytometry Core facility using a BD 5 laser SORP FACS Vantage SE Diva system or Facsaria (BD Biosciences). FACS data and ∑Median statistics were analyzed using FlowJo 10.8.1 software (Tree Star, Inc.).

## References

1. MacKenzie G, Maguire J. The role of ovarian hormone-derived neurosteroids on the regulation of GABAA receptors in affective disorders. Psychopharmacology (Berl). 2014;231(17):3333–42. Epub 2014/01/10. doi: 10.1007/s00213-013-3423-z. PMID: 24402140.

2. Maguire JL, Stell BM, Rafizadeh M, Mody I. Ovarian cycle-linked changes in GABA(A) receptors mediating tonic inhibition alter seizure susceptibility and anxiety. Nat Neurosci. 2005;8(6):797–804. Epub 20050515. doi: 10.1038/nn1469. PMID: 15895085.

3. Rogawski MA, Loya CM, Reddy K, Zolkowska D, Lossin C. Neuroactive steroids for the treatment of status epilepticus. Epilepsia. 2013;54 Suppl 6(0 6):93–8. doi: 10.1111/epi.12289. PMID: 24001085.

4. Blanco MJ, La D, Coughlin Q, Newman CA, Griffin AM, Harrison BL, Salituro FG. Breakthroughs in neuroactive steroid drug discovery. Bioorg Med Chem Lett. 2018;28(2):61–70. Epub 20171202. doi: 10.1016/j.bmcl.2017.11.043. PMID: 29223589.

5. Maguire JL, Mennerick S. Neurosteroids: mechanistic considerations and clinical prospects. Neuropsychopharmacology. 2024;49(1):73–82. Epub 20230627. doi: 10.1038/s41386-023-01626-z. PMID: 37369775.

6. Clayton AH, Lasser R, Parikh SV, Iosifescu DV, Jung J, Kotecha M, Forrestal F, Jonas J, Kanes SJ, Doherty J. Zuranolone for the Treatment of Adults With Major Depressive Disorder: A Randomized, Placebo-Controlled Phase 3 Trial. Am J Psychiatry. 2023;180(9):676–84. Epub 20230503. doi: 10.1176/appi.ajp.20220459. PMID: 37132201.

7. Singhal M, Modi N, Bansal L, Abraham J, Mehta I, Ravi A. The Emerging Role of Neurosteroids: Novel Drugs Brexanalone, Sepranolone, Zuranolone, and Ganaxolone in Mood and Neurological Disorders. Cureus. 2024;16(7):e65866. Epub 20240731. doi: 10.7759/cureus.65866. PMID: 39219949.

8. Longone P, Rupprecht R, Manieri GA, Bernardi G, Romeo E, Pasini A. The complex roles of neurosteroids in depression and anxiety disorders. Neurochem Int. 2008;52(4-5):596–601. Epub 20071006. doi: 10.1016/j.neuint.2007.10.001. PMID: 17996986.

9. Mishra S, Elliott H, Marwaha R. Premenstrual Dysphoric Disorder. StatPearls. Treasure Island (FL) 2025.

10. Backstrom T, Bixo M, Johansson M, Nyberg S, Ossewaarde L, Ragagnin G, Savic I, Stromberg J, Timby E, van Broekhoven F, van Wingen G. Allopregnanolone and mood disorders. Prog Neurobiol. 2014;113:88–94. Epub 20130823. doi: 10.1016/j.pneurobio.2013.07.005. PMID: 23978486.

11. Hantsoo L, Epperson CN. Allopregnanolone in premenstrual dysphoric disorder (PMDD): Evidence for dysregulated sensitivity to GABA-A receptor modulating neuroactive steroids across the menstrual cycle. Neurobiol Stress. 2020;12:100213. Epub 20200204. doi: 10.1016/j.ynstr.2020.100213. PMID: 32435664.

12. Backstrom T, Das R, Bixo M. Positive GABA(A) receptor modulating steroids and their antagonists: Implications for clinical treatments. J Neuroendocrinol. 2022;34(2):e13013. Epub 20210801. doi: 10.1111/jne.13013. PMID: 34337790.

13. Bixo M, Ekberg K, Poromaa IS, Hirschberg AL, Jonasson AF, Andreen L, Timby E, Wulff M, Ehrenborg A, Backstrom T. Treatment of premenstrual dysphoric disorder with the GABA(A) receptor modulating steroid antagonist Sepranolone (UC1010)-A randomized controlled trial. Psychoneuroendocrinology. 2017;80:46–55. Epub 20170301. doi: 10.1016/j.psyneuen.2017.02.031. PMID: 28319848.

14. Thompson SM. Modulators of GABA(A) receptor-mediated inhibition in the treatment of neuropsychiatric disorders: past, present, and future. Neuropsychopharmacology. 2024;49(1):83–95. Epub 20230914. doi: 10.1038/s41386-023-01728-8. PMID: 37709943.

15. Backstrom T, Doverskog M, Blackburn TP, Scharschmidt BF, Felipo V. Allopregnanolone and its antagonist modulate neuroinflammation and neurological impairment. Neurosci Biobehav Rev. 2024;161:105668. Epub 20240410. doi: 10.1016/j.neubiorev.2024.105668. PMID: 38608826.

16. Llansola M, Mincheva G, Arenas YM, Izquierdo-Altarejos P, Pedrosa MA, Blackburn TP, Backstrom T, Scharschmidt BF, Doverskog M, Felipo V. Golexanolone Attenuates Neuroinflammation, Fatigue, and Cognitive and Motor Impairment in Diverse Neuroinflammatory Disorders. Pharmaceuticals (Basel). 2025;18(11). Epub 20251118. doi: 10.3390/ph18111757. PMID: 41304999.

17. Yilmaz C, Karali K, Fodelianaki G, Gravanis A, Chavakis T, Charalampopoulos I, Alexaki VI. Neurosteroids as regulators of neuroinflammation. Front Neuroendocrinol. 2019;55:100788. Epub 20190909. doi: 10.1016/j.yfrne.2019.100788. PMID: 31513776.

18. Stell BM, Brickley SG, Tang CY, Farrant M, Mody I. Neuroactive steroids reduce neuronal excitability by selectively enhancing tonic inhibition mediated by delta subunit-containing GABAA receptors. Proc Natl Acad Sci U S A. 2003;100(24):14439–44. Epub 2003/11/19. doi: 10.1073/pnas.2435457100. PMID: 14623958.

19. Wohlfarth KM, Bianchi MT, Macdonald RL. Enhanced neurosteroid potentiation of ternary GABA(A) receptors containing the delta subunit. J Neurosci. 2002;22(5):1541–9. doi: 10.1523/JNEUROSCI.22-05-01541.2002. PMID: 11880484.

20. Kaur KH, Baur R, Sigel E. Unanticipated structural and functional properties of delta-subunit-containing GABAA receptors. J Biol Chem. 2009;284(12):7889–96. Epub 20090113. doi: 10.1074/jbc.M806484200. PMID: 19141615.

21. Eaton MM, Bracamontes J, Shu HJ, Li P, Mennerick S, Steinbach JH, Akk G. gamma-aminobutyric acid type A alpha4, beta2, and delta subunits assemble to produce more than one functionally distinct receptor type. Mol Pharmacol. 2014;86(6):647–56. Epub 2014/09/23. doi: 10.1124/mol.114.094813. PMID: 25238745.

22. Wongsamitkul N, Baur R, Sigel E. Toward Understanding Functional Properties and Subunit Arrangement of alpha4beta2delta gamma-Aminobutyric Acid, Type A (GABAA) Receptors. J Biol Chem. 2016;291(35):18474–83. Epub 2016/07/07. doi: 10.1074/jbc.M116.738906. PMID: 27382064.

23. Sente A, Desai R, Naydenova K, Malinauskas T, Jounaidi Y, Miehling J, Zhou X, Masiulis S, Hardwick SW, Chirgadze DY, Miller KW, Aricescu AR. Differential assembly diversifies GABAA receptor structures and signalling. Nature. 2022;604(7904):190–4. Epub 2022/04/01. doi: 10.1038/s41586-022-04517-3. PMID: 35355020.

24. Botzolakis EJ, Gurba KN, Lagrange AH, Feng HJ, Stanic AK, Hu N, Macdonald RL. Comparison of gamma-Aminobutyric Acid, Type A (GABAA), Receptor alphabetagamma and alphabetadelta Expression Using Flow Cytometry and Electrophysiology: Evidence for alternative subunit stoichiometries and arrangements. J Biol Chem. 2016;291(39):20440–61. Epub 2016/08/06. doi: 10.1074/jbc.M115.698860. PMID: 27493204.

25. Feng HJ, Forman SA. Comparison of alphabetadelta and alphabetagamma GABAA receptors: Allosteric modulation and identification of subunit arrangement by site-selective general anesthetics. Pharmacol Res. 2018;133:289–300. Epub 2018/01/03. doi: 10.1016/j.phrs.2017.12.031. PMID: 29294355.

26. Liao VWY, Chebib M, Ahring PK. Efficient expression of concatenated alpha1beta2delta and alpha1beta3delta GABAA receptors, their pharmacology and stoichiometry. Br J Pharmacol. 2021;178(7):1556–73. Epub 2021/01/26. doi: 10.1111/bph.15380. PMID: 33491192.

27. Koinas D, Zhou X, Wu B, Miller KW, Bruzik KS. Novel Spiro-Barbiturates Can Reverse the Action of General Anesthetics on the GABA(A)R. J Med Chem. 2025;68(8):8025–45. Epub 20250407. doi: 10.1021/acs.jmedchem.4c02514. PMID: 40193703.

28. Koinas D, Zhou X, Wu B, Bruzik KS, Miller KW. Spiro Hydantoins Can Reverse the Action of Positive Allosteric Modulators on GABAARs. ACS Med Chem Lett. 2025;16:2078–83. Epub 06 May 2025. doi: 10.1021/acsmedchemlett.5c00499.

29. Miehling J. Mechanism of anaesthetic activation, combination and antagonism [Thesis]. Apollo–University of Cambridge: University of Cambridge; 2022. 10.17863/CAM.87926

30. Zuo Y, Zhao Y, Liu G, Sun Q. Recent advances in GABA(A) receptor targeting ligands. Eur J Med Chem. 2026;308:118651. Epub 20260205. doi: 10.1016/j.ejmech.2026.118651. PMID: 41719803.

31. Stromberg J, Lundgren P, Taube M, Backstrom T, Wang M, Haage D. The effect of the neuroactive steroid 5beta-pregnane-3beta, 20(R)-diol on the time course of GABA evoked currents is different to that of pregnenolone sulphate. Eur J Pharmacol. 2009;605(1-3):78–86. Epub 20090110. doi: 10.1016/j.ejphar.2008.12.038. PMID: 19168059.

32. Chang Y, Ghansah E, Chen Y, Ye J, Weiss DS. Desensitization mechanism of GABA receptors revealed by single oocyte binding and receptor function.[erratum appears in J Neurosci 2002 Oct 15;22(20):1b Note: Chang YongChang [corrected to Chang Yongchang]]. Journal of Neuroscience. 2002;22(18):7982–90.

33. Germann AL, Pierce SR, Tateiwa H, Sugasawa Y, Reichert DE, Evers AS, Steinbach JH, Akk G. Intrasubunit and Intersubunit Steroid Binding Sites Independently and Additively Mediate alpha1beta2gamma2L GABA(A) Receptor Potentiation by the Endogenous Neurosteroid Allopregnanolone. Mol Pharmacol. 2021;100(1):19–31. Epub 20210506. doi: 10.1124/molpharm.121.000268. PMID: 33958479.

34. Legesse DH, Fan C, Teng J, Zhuang Y, Howard RJ, Noviello CM, Lindahl E, Hibbs RE. Structural insights into opposing actions of neurosteroids on GABA(A) receptors. Nat Commun. 2023;14(1):5091. Epub 20230822. doi: 10.1038/s41467-023-40800-1. PMID: 37607940.

35. Dostalova Z, Liu A, Zhou X, Farmer SL, Krenzel ES, Arevalo E, Desai R, Feinberg-Zadek PL, Davies PA, Yamodo IH, Forman SA, Miller KW. High-level expression and purification of Cys-loop ligand-gated ion channels in a tetracycline-inducible stable mammalian cell line: GABAA and serotonin receptors. Protein Sci. 2010;19(9):1728–38. Epub 2010/07/28. doi: 10.1002/pro.456. PMID: 20662008.

36. Dostalova Z, Zhou X, Liu A, Zhang X, Zhang Y, Desai R, Forman SA, Miller KW. Human alpha1beta3gamma2L gamma-aminobutyric acid type A receptors: High-level production and purification in a functional state. Protein Sci. 2014;23(2):157–66. Epub 2013/11/30. doi: 10.1002/pro.2401. PMID: 24288268.

37. Zhou X, Desai R, Zhang Y, Stec WJ, Miller KW, Jounaidi Y. High-level production and purification in a functional state of an extrasynaptic gamma-aminobutyric acid type A receptor containing alpha4beta3delta subunits. PLoS One. 2018;13(1):e0191583. Epub 2018/01/21. doi: 10.1371/journal.pone.0191583. PMID: 29352320.

38. Jensen ML, Wafford KA, Brown AR, Belelli D, Lambert JJ, Mirza NR. A study of subunit selectivity, mechanism and site of action of the delta selective compound 2 (DS2) at human recombinant and rodent native GABA(A) receptors. Br J Pharmacol. 2013;168(5):1118–32. Epub 2012/10/16. doi: 10.1111/bph.12001. PMID: 23061935.

